# *slim shady* is a novel allele of PHYTOCHROME B present in the T-DNA line SALK_015201

**DOI:** 10.1101/2021.02.12.430994

**Authors:** Linkan Dash, Robert E. McEwan, Christian Montes, Ludvin Mejia, Justin W. Walley, Brian P. Dilkes, Dior R. Kelley

## Abstract

Auxin is a hormone that is required for hypocotyl elongation during seedling development. In response to auxin rapid changes in transcript and protein abundance occur in hypocotyls and some auxin responsive gene expression is linked to hypocotyl growth. To functionally validate proteomic studies, a reverse genetics screen was performed on mutants in auxin-regulated proteins to identify novel regulators of plant growth. This uncovered a long hypocotyl mutant, which we called *slim shady*, in an annotated insertion line in *IMMUNOREGULATORY RNA-BINDING PROTEIN (IRR*). Overexpression of the *IRR* gene failed to rescue the *slim shady* phenotype and characterization of a second T-DNA allele of *IRR* found that it had a wild-type hypocotyl length. The *slim shady* mutant has an elevated expression of numerous genes associated with the brassinosteroid-auxin-phytochrome (BAP) regulatory module compared to wild-type, including transcription factors that regulate brassinosteroid, auxin and phytochrome pathways. Additionally, *slim shady* seedlings fail to exhibit a strong transcriptional response to auxin. Using whole genome sequence and transcriptomics data for SALK_015201C we determined that a novel single nucleotide polymorphism in *PHYTOCHROME B* was responsible for the *slim shady* phenotype. This is predicted to convert induce a frameshift and premature stop codon at leucine 1125, within the histidine kinase-related domain of the carboxy terminus of PHYB, which is required for phytochrome signaling and function. Genetic complementation analyses with *phyb-9* confirmed that slim shady is a mutant allele of *PHYB*. This study advances our understanding of the molecular mechanisms in seedling development, by furthering our understanding of how light signaling is linked to auxin dependent cell elongation. Furthermore, this study highlights the importance of confirming the genetic identity of research material before attributing phenotypes to known mutations sourced from T-DNA stocks.

## Introduction

In plants, phytohormones work in concert to coordinate cell divisions, cell expansions and drive morphogenesis. Hypocotyl development in Arabidopsis has been an effective model system for studying the interaction between phytohormones and their roles in responses to various environmental signals. Arabidopsis hypocotyl has approximately 20 cell files from apex to base that do not proliferate post embryonically and their elongation largely depends on cell expansion (Gendreau et al., 1997). Consequently, post-embryonic cell expansion determines hypocotyl elongation in Arabidopsis and is largely influenced by internal factors like hormones and external environmental factors like light and temperature.

Phytochromes are light sensing proteins that dynamically exist in two interconvertible isoforms, the red light absorbing Pr form and the far-red light absorbing Pfr form that is considered to be biologically active (Rockwell et al., 2006). In light conditions, cytosolic biologically-inactive Pr form absorbs red light and undergoes rapid conformational change to the Pfr form and is imported to the nucleus (Nagatani, 2004). Nuclear localized Pfr dimers physically interact with PHYTOCHROME INTERACTING FACTORS (PIFs), a group of basic helix-loop-helix transcription factors that bind to ‘G-box’ elements in promoters to facilitate transcription (Leivar and Monte, 2014; Leivar et al., 2012).

Light activated phytochromes phosphorylates the PIFs and trigger their 26S proteasome-mediated degradation and removing the from promoters (Al-Sady et al., 2006; Park et al., 2012). The PIFs can directly trigger expression of downstream genes that can increase auxin biosynthesis and/or signaling (Franklin et al., 2011; Nozue et al., 2011; Hornitschek et al., 2012; Sun et al., 2012; Li et al., 2012b; Goyal et al., 2016). The PIFs also regulate both brassinosteroid and auxin responses by physically interacting with their respective regulatory transcription factors, BRASSINAZOLE RESISTANT1 (BZR1) and AUXIN RESPONSE FACTORs (ARFs) (Oh et al., 2014).

The central signaling network that positively regulates hypocotyl growth in Arabidopsis consists of three major families of interdependently working transcription factors-BRASSINAZOLE RESISTANT1 (BZR1), AUXIN RESPONSE FACTORs (ARFs) and the PHYTOCHROME INTERACTING FACTORs (PIFs), collectively referred to as the BZR/ARF/PIF module or the ‘BAP module’ (Bouré et al., 2019). The BAP module works in concert to activate downstream genes that promote cell elongation in the hypocotyl (Favero, 2020). Plant hormone gibberellins also contribute towards hypocotyl elongation by modulating PIF activity and stability (De Lucas et al., 2008; Feng et al., 2008; Li et al., 2016). While auxin can rapidly regulate gene expression at the transcript and protein level (Bargmann et al., 2014; Nemhauser et al., 2006; Clark et al., 2019; Pu et al., 2019).

T-DNA mutants are a valuable resource to the Arabidopsis community and have greatly facilitated genetic screens and functional genomics (Dilkes and Feldmann, 1998; O’Malley and Ecker, 2010; Page and Grossniklaus, 2002). But they are not without their drawbacks. Additional mutant alleles unassociated with T-DNA insertions (also called untagged T-DNA mutants) have been reported on numerous occasions and can be associated with duplications/translocations, insertion/deletions or other complex genetic changes (Tax and Vernon, 2001). Notable examples include second-site mutations in the abp1-5 and phyb-9 backgrounds (Enders et al., 2015; Gao et al., 2015; Yoshida et al., 2018), a new set of transparent testa alleles (Jiang et al., 2020), and a mutation in nrpd1a-3 associated with ROOT HAIR DEFECTIVE 6 (RHD6) (David et al., 2019).

In this study we performed a reverse genetic screen based on our auxin responsive proteomics data (Clark et al., 2019) and publicly available Arabidopsis T-DNA insertion alleles (O’Malley and Ecker, 2010; Alonso et al., 2003). From this screen we identified a T-DNA line with a long hypocotyl phenotype which we initially called slim shady and was recently published as IMMUNOREGULATORY RNA BINDING PROTEIN (IRR) (Dressano et al., 2020). A long hypocotyl phenotype in SALK_015201 lines had been previously reported (Petrov et al., 2013) but a second T-DNA allele in IRR had a wild-type phenotype, prompting us to reconsider the genetic lesion responsible for the long hypocotyl phenotype. Analysis of whole genome sequence data available for this SALK line (Hu et al., 2019) identified a single base pair deletion within the coding sequence of PHYB. Transcriptomic comparison of SALK_015201 and wild type seedlings with and without auxin determined that genes associated with the BAP regulatory module (Bouré et al., 2019) were altered in this T-DNA background. Genetic complementation analyses confirmed the presence of a new phyb allele, which we call slim shady, in SALK_015201. This mutation is predicted to generate a truncated PHYB protein. This study serves as a reminder for the Arabidopsis community when working with T-DNA alleles to backcross material, seek multiple alleles, and remain vigilant about the high rate of loss of function mutations in these lines.

## Materials and Methods

### Plant Material

All seed stocks used in this study were obtained from the Arabidopsis Biological Resource Center (ABRC) at Ohio State University. Arabidopsis thaliana plants used in this study were Columbia (Col-0) ecotype. SALK_015201 encodes the *irr-1* knock-out allele (also annotated as SALK_015201C and SALKseq_015201) and SALK_066572 encodes the *irr-2* knock-out allele which were previously characterized (Dressano et al., 2020; Petrov et al., 2013). Homozygous *phyB-9* seed stocks (ABRC stock number CS6217) were also previously characterized (Reed et al., 1994).

For transcriptomic profiling, Col-0 and homozygous SALK_015201C seeds were surface sterilized using 50% bleach and 0.01% Triton X-100 for 10 minutes and then washed five times with sterile water. Seeds were then imbibed in sterile water for two days at 4°C and then transferred to 0.5X Linsmaier-Skoog (LS) medium plates solidified with 0.8% agar and overlaid with sterile 100 micron nylon mesh squares to facilitate tissue harvesting. Seedlings were grown under long day photoperiods (16 h light/8 h dark) at 23°C. Sixdayold seedlings were treated with 1 indole-3-acetic acid (IAA) dissolved in 95% ethanol (“auxin”) or an equivalent volume of 95% ethanol (“mock”) for 24 hours by transferring the seedlings on mesh squares to square petri dishes containing 10 mL of fresh 0.5X LS supplemented with IAA or mock solvent for the specified time. Following treatments, the seedlings were then harvested, weighed, and immediately frozen in liquid nitrogen; approximately 1 g of seedling tissue was collected per replicate/genotype. Four independent biological replicates were generated for each genotype and treatment.

For phenotyping assays, seeds were surface sterilized using 50% bleach and 0.01% Triton X-100 for 10 minutes and then washed five times with sterile water. Seeds were then imbibed in sterile water for two days at 4°C and then transferred to 0.5X LS medium plates. Seedlings were grown under long day photoperiods (16 h light/8 h dark) and dark (24 h dark post 2 h light incubation) at 23°C. For auxin response assays, five-day-old seedlings were transferred to either 0.5X LS plates or 0.5X LS plates supplemented with 1 IAA and grown for another two days.

A 35S:GFP-IRR transgenic line was created using Agrobacterium mediated transformation into the SALK_015201C background. Sevendayold Col-0 seedlings were snap frozen in liquid nitrogen followed by tissue lysis using mortar and pestle. Extraction of total RNA from the harvested tissue was done using TRIzol™ Reagent (Invitrogen, catalog number-15596026) followed by purification using RNeasy^®^ mini-kit (Qiagen, catalog number-74106). The purified total RNA was used to synthesize cDNA using SuperScript™ IV Reverse Transcriptase kit from Invitrogen (catalog number-18090010). The synthesized total cDNA was used as a template for PCR amplifying the full-length *IRR* cDNA, using primers modified for directional cloning (supplemental table 2). The blunt end PCR product was cloned into the pENTR™/D-TOPO^®^ donor vector, using the gateway cloning kit provided by Invitrogen (catalog number: K240020). The entry clone was then transformed into chemically competent DH5cells. Transformants were selected for kanamycin resistance and tested by colony PCR and restriction digest. The cloned plasmids were then isolated from their bacterial cultures using GeneJET Plasmid Miniprep Kit from Thermo Fisher scientific (catalog number-K0503).

In the next step, an LR reaction was set between the entry clone and the gateway destination vector pGWB606 (Nakamura et al., 2010) using LR clonase™ enzyme mix provided by Invitrogen (catalog number-11791020). In this case the transformants in *E. coli* were selected by spectinomycin resistance, colony PCR with primers specific to the chimeric *GFP:IRR cDNA* (supplemental table 2). The final clone was further confirmed by sanger sequencing (data not shown), followed by their transformation into chemically competent *Agrobacterium* cells of GV3101 strain.

Columbia-0 and SALK_015201 plants were grown under long day photo period (16 hr light, 8 hr dark) until they reached bolting. Floral dip plant transformation protocol was performed on these plants and they were left to grow normally until senescence. Harvested seeds from the dipped plants were sown on soil and were allowed to grow normally until 2-4 leaves stage. Juvenile plants were sprayed with a 1:1000 dilution of 120mg/ml BASTA (Finale) for selecting the positive transformants which were further confirmed through genotyping for the presence of chimeric *GFP-IRR* gene (data not shown).

### Genotyping

All primers used for genotyping are provided in Supplemental Table 2. Primers to genotype SALK alleles were designed using the SALK T-DNA verification primer design tool at the SALK SiGNAL website (http://signal.salk.edu/tdnaprimers.2.html). A derived cleaved amplified polymorphism (dCAPS) marker was designed to differentiate between the WT and slim shady alleles of PHYB using the dCAPS 2.0 finder tool (Bui and Liu, 2009). For the dCAPS analysis, a 280 bp region of *PHYB* was amplified using the dCAPS primers (Supplemental Table 2) with DreamTaq polymerase (Thermo Fisher Scientific) and subsequently digested with ApoI-HF (New England Biolabs) according to manufacturer’s protocols. The digested PCR products were analyzed on a 4% agarose gel. The WT allele of *PHYB* is expected to be cleaved by ApoI-HF while the mutant *slim shady* allele is not cleaved.

### Genetic Analyses

Homozygous single mutants were crossed to generate F1 seedlings for complementation analyses: SALK_015201C was crossed to both Col-0 and *phyb-9*. As a control, *irr-2* was also crossed to *phyb-9*. The resulting F1 seeds were surface sterilized as described above and plated on 0.5X LS for phenotyping.

### Transcriptomic and GO Enrichment Analyses

RNA was extracted from 0.1 g of sevendayold seedlings (Col-0 and SALK_015201) using Trizol followed by column clean up using the Quick-RNA plant kit (Zymo). Total RNA concentrations were determined using a NanoDrop and Qubit. RNA quality was checked via Bioanalyzer at the ISU DNA Facility. QuantSeq 3’ mRNA libraries were prepared using the Lexogen 3’ mRNA-seq FWD kit and sequenced on an Illumina HiSeq 3000 as 50 bp reads at the ISU DNA Facility. QuantSeq reads were mapped to the TAIR10 genome and differential gene expression analysis was performed using PoissonSeq implemented in R (Li et al., 2012a). Transcripts with a FDR cutoff of 0.05 and a log fold change 1.75 were defined as high confidence differentially expressed. Gene Ontology (GO) enrichment analyses were performed using BiNGO in Cytoscape (Maere et al., 2005) with either upregulated or downregulated genes as the input.

### SNPs Calling and Informatics

DNA polymorphisms were identified following alignment of whole-genome sequences to the *A. thaliana* TAIR10 genome. Genomic sequence data for A. thaliana accessions SALK_015201 and CS85255 were retrieved from the Sequence Read Archive (SRA) at the National Center for Biotechnology Information (NCBI) with IDs SRR5249176 and SRR5249156 respectively (Hu et al., 2019). These paired-end 150-bp reads were mapped to the TAIR10 reference genome (Lamesch et al., 2012) using BWA-MEM (version 0.7.17) (Li and Durbin, 2009). Duplicate reads were identified and removed from alignment files using the *rmdup* command of SAMtools (version 0.1.18) (Li and Durbin, 2009).

SNP and small indel calling was carried out as a three-step process. First, the SAMtools *mpileup* command was used to transpose mapped reads to the reference genome and compute genotype likelihood. Next, the BCFtools *view* command was used to perform actual variant calling, and finally, preliminary quality filtering was done with *varFilter* command of the *vcfutil.pl* script with the “-D100” option to exclude variants with more than 100 reads coverage. Further filtering was performed with SnpSift (version 3.6) (Cingolani et al., 2012) to keep only variants that have a phred-scale quality of at least 20. The VCFtools *exclude-positions* command was used to remove false-positive variants using a list of variants common to phenotypically unaffected individuals. These variants are deemed false-positives because they occur in numerous lineages and thus cannot be causal for a phenotype seen only in a single lineage (Addo-Quaye et al., 2017, 2018). SnpEff (version 3.6) (Cingolani et al., 2012) was used with TAIR10 gene annotation to predict the effect of the remaining variants on gene function.

To check for incomplete integration by the T-DNA or Ti plasmids and confirm the identities of the SRA samples, sequences for pDs-Lox, pBIN-pROK2 and M11311.1 Ti plasmid were added as extra chromosomes to the TAIR10 reference genome to create a new TAIR10 pseudo-reference genome that could detect these T-DNA sequences by alignment. Reads from SRR5249176 and SRR5249156 were independently aligned to the pseudo-reference with BWA-MEM. Alignment files were converted from SAM to BAM format with the SAMtools *view* command prior to duplicate read removal with the *rmdup* command. Deduplicated BAM files for both samples were then viewed in IGV to check for the presence of Ds-Lox and pBIN-pROK2 T-DNA.

### RNA polymorphism calling by alignment to *A. thaliana*

RNA-seq reads for twelve individuals comprising Pep-treated and untreated *irr-1* mutant and wild-type plants (three biological replicates per genotype per condition) were obtained from the NCBI SRA with IDs SRR11218900 - SRR11218911 (Dressano et al., 2020). The paired-end 150-bp reads were aligned to the TAIR10 reference genome (Lamesch et al., 2012) using the spliced alignment option of the STAR aligner (Dobin et al., 2013). The output was processed for duplicate read removal with SAMtools *rmdup* and passed to the variant calling pipeline described above, with the notable exception of the exclusion of a false-positive variant removal step.

Aligned reads were additionally analyzed to identify differentially expressed genes (DEGs) induced by Pep treatment using DESeq2 (Love et al., 2014). The *htseq-count* function of HTSeq (Anders and Huber, 2010) was used to project alignment files to TAIR10.32 gene/transcript annotation to count reads/fragments to produce gene level count matrices used as input for DESeq2. Genes were considered differentially expressed between Pep treated vs untreated if they had an adjusted p value 0.1 (i.e., false discovery rate of 0.1 was applied).

### Hypocotyl and Root Phenotyping

Intact seedlings (seven day-old) were imaged on a flat-bed scanner (Epson V600). Measurements of hypocotyl and primary root lengths were performed using ImageJ. One-way Analysis of Variance (ANOVA) was performed for comparing the F1 and single mutant genotypes against control Col-0. Here the hypocotyl length was taken as the dependent variable with genotype of the plants as the only independent variable.

### Sanger Sequencing and Sequence Alignments

A 644 bp region of *PHYB* was amplified from Col-0, SALK_015201 and *irr-2* genomic DNA using gene specific primers (Supplemental Table 2) with Phusion polymerase (Thermo Fisher Scientific) and cleaned up using the GeneJET PCR Purification Kit from Thermofisher (catalog number K0701). Purified PCR products were subjected to standard Sanger sequencing at the Iowa State University DNA Facility on an Applied Biosystems 3730xl DNA Analyzer. Sequence files were trimmed in SnapGene and aligned using CLUSTAL Omega.

### Accession Numbers

Sequence data from this article can be found at the NCBI BioProject database under SubmissionID: SUB8925253 and BioProject ID: PRJNA694682. The project information will be accessible with the following link within a few days of the release date upon publication: http://www.ncbi.nlm.nih.gov/bioproject/694682. Data analyses performed herein utilized public repository data from Hu et. al. 2019 *BMC Bioinformatics* deposited at the NCBI SRA database under accession SRR5249176 and (Dressano et al., 2020) deposited at NCBI GEO GSE146282.

## Results

### SALK_015201 (*irr-1*) displays a long hypocotyl phenotype

Auxin regulated gene expression has been well-characterized and influences many aspects of seedling growth and development. We recently quantified auxin regulated proteome remodeling in five-day-old Arabidopsis seedlings (Clark et al., 2019; Kelley et al., 2017) and found widespread changes in protein abundance not linked to transcriptional changes. In order to explore and test this idea further we examined auxin responsive proteins from our dataset to identify putative candidate proteins which could underlie post-transcriptional auxin-mediated gene expression. One top candidate was the protein IMMUNOREGULATORY RNA-BINDING PROTEIN (IRR; AT3G23900) which increased in abundance in response to auxin (Kelley et al., 2017) and is known to influence mRNA splicing in Arabidopsis (Dressano et al., 2020).

In order to characterize loss-of-function of *IRR* and determine if *IRR* is required for auxin regulated gene expression, we obtained two publicly available SALK T-DNA lines in this gene (Figure 1). Initial phenotyping of the SALK_015201C line indicated that this T-DNA line exhibited a long hypocotyl phenotype (Figure 1), which had been previously reported (Petrov et al., 2013). Conversely, the other SALK allele annotated in the *IRR* gene, SALK_066572 (*irr-2*) had a wild-type seedling phenotype. Based on our initial phenotypic characterization we termed the SALK_015201C mutant *slim shady*. Notably fifteen-day-old *irr-1* and *irr-2* roots have been described as longer compared to WT (Dressano et al., 2020), but we did not observe any differences in primary root length in either SALK allele compared to WT (data not shown). This could be due to different growth media conditions and/or developmental age examined between the studies. Homozygous seed stocks of SALK_015201C and SALK_066572 were confirmed using gene specific and T-DNA primers listed in Supplemental Table 2.

**Fig. 1.**
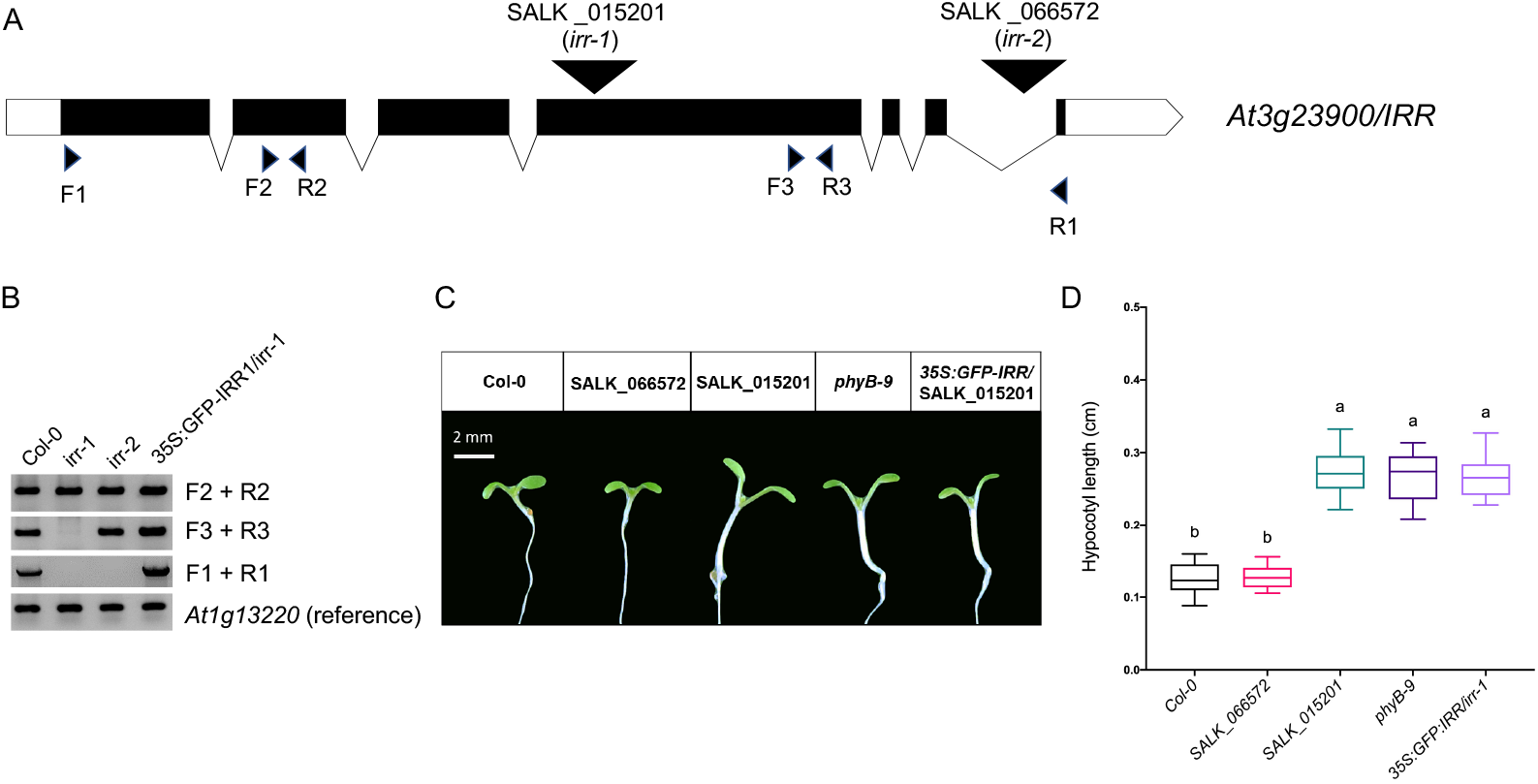
(A) Annotated T-DNA alleles of *IMMUNOREGULATORY RNA BINDING PROTEIN (IRR*) include SALK_015201 (*irr*-1) and SALK_066572 (*irr*-2). Exons are indicated as black bars, introns as lines, untranslated regions (UTR) as white boxes and T-DNA insertions as inverted triangles. (B) RT-PCR of these alleles indicates that both *irr-1* and *irr-2* are null alleles of *IRR* that produce upstream truncated transcripts, while a *35S::GFP-IRR* transgenic line in the SALK_015201 background has restored expression of IRR. *At1g13220* was used as a control gene. (C) Sevendayold *phyb-9*, SALK_015201 and *35S:GFP-IRR/irr-1* seedlings display long hypocotyls compared to Col-0 and *irr-2*. Scale bar = 2 mm. (D) Quantification of hypocotyl lengths. Letters a and b indicate the significant statistical differences between the hypocotyl lengths of different genotypes, determined by a oneway ANOVA followed by Tukey’s HSD post hoc test with p 0.05.

**Fig. 2.**
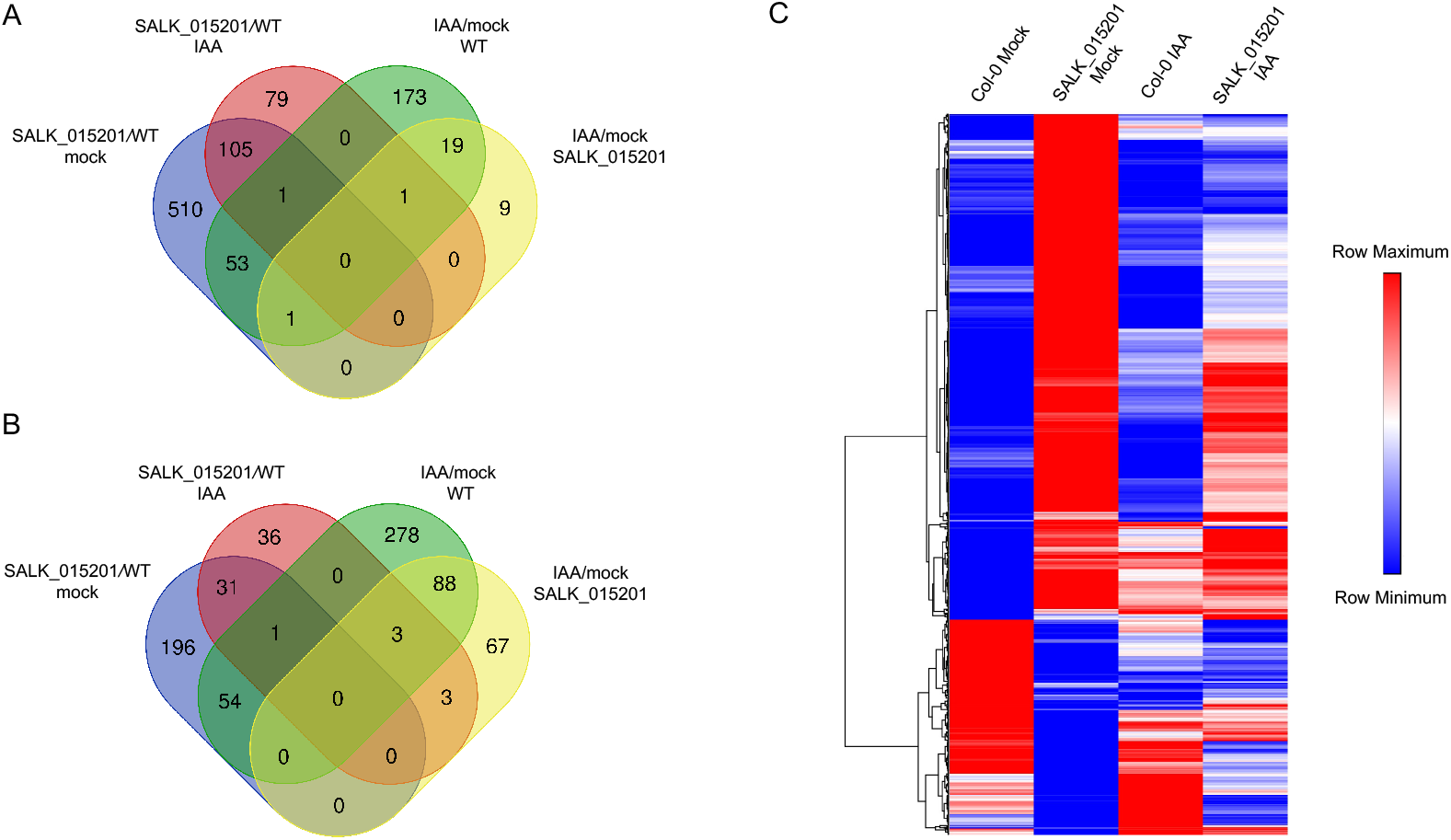
Differential gene expression (DEG) analysis in SALK_015201 compared to Col-0. Venn diagrams of (A) upregulated genes and (B) downregulated genes. (C) Heat map of all 952 DEGs depicting their mean normalized read counts where the scale bar ranges from lowest (blue) to highest (red) relative expression value within a row generated by hierarchical clustering using one-minus Pearson correlation matrix.

To characterize *IRR* gene expression in SALK_015201C (*irr-1*) and SALK_066572 (irr-2) we performed reverse transcription polymerase chain reaction (RT-PCR) analysis using gene specific primers (Supplemental Table 2) designed to amplify either full-length transcript or truncated mRNA amplicons both up- and downstream of the T-DNA insertion sites (Figure 1A). *At1g13220* was used as a reference gene for normalization (Czechowski, 2005).

Consistent with a previous report, neither allele produced a full-length *IRR* transcript, but truncated upstream transcripts were detected in both alleles (Figure 1B). *IRR* expression was examined using primers that amplified target regions in exon 2 (upstream of the annotated T-DNA insertion site), exon 4 (downstream of the annotated T-DNA insertion sites) as well as the full length of *IRR* cDNA. The expression of a *35S:GFP-IRR* transgene in the SALK_015201 background restored expression of *IRR* back to normal (Figure 1B).

Loss of phytochrome function is classically associated with a long hypocotyl phenotype (Franklin and Quail, 2010). As a phenotypic comparison we quantified the hypocotyl lengths of the SALK alleles and the *35S:GFP-IRR* line relative to WT and *phyb-9* (Figure 1C). The SALK_015201 line exhibits an elongated hypocotyl phenotype as compared to the Col-0 and SALK_066572 (irr-2) (Figure 1D). While the hypocotyl length of 7 day-old SALK_015201 seedlings were found to be statistically indifferent from *phyB-9* (Figure 1D). Transgenic lines expressing *35S:GFP-IRR* failed to restore the hypocotyl phenotype in SALK_015201 back to normal (Figure 1C, D).

### BAP module gene expression is elevated in SALK_015201 seedlings

We hypothesized that the long hypocotyl phenotype observed in SALK_015201 seedlings may be due to altered auxin responses and/or light signaling. To test this hypothesis, we performed RT-qPCR on three key marker genes, *INDOLE-3-ACETIC ACID INDUCIBLE 14/SOLITARY ROOT (IAA14/SLR), PHYTOCHROME A (PHYA*) and *PHYB* (Supplemental Figure 1). All three genes show elevated expression in SALK_015201 compared to WT. We then performed a transcriptomic analysis to determine global gene expression patterns in SALK_015201 both in the presence and absence of auxin treatment using the 3’ end mRNA sequencing method (Moll et al., 2014).

Differential gene expression (DE) analysis was determined using the PoissonSeq package implemented in R (Li et al., 2012a). We applied a FDR threshold of 0.05 and a log fold change between comparisons (i.e. WT versus SALK_015201 mock treated) 1.75 to define DE genes. We found 670 genes upregulated and 282 genes to be downregulated in SALK_015201 compared to WT (Supplemental Table 1). In response to a 24-hour auxin treatment (1 indole-3-acetic acid) we observed 186 genes upregulated and 74 genes to be down-regulated in SALK_015201 compared to WT (Supplemental Table 1). Gene Ontology (GO) terms such as auxin response, light response (blue, red, and far-red), cell wall modification and shade avoidance response were enriched for the set of highly expressed genes in SALK_015201 as compared to the WT (Figure 3 and Supplemental Figure 2). Downregulated genes were enriched in GO terms such as nuclear mRNA splicing via. Spliceosome, Glucosinolate biosynthetic process, Response to Jasmonic acid stimulus (Supplemental Figure 3).

**Fig. 3.**
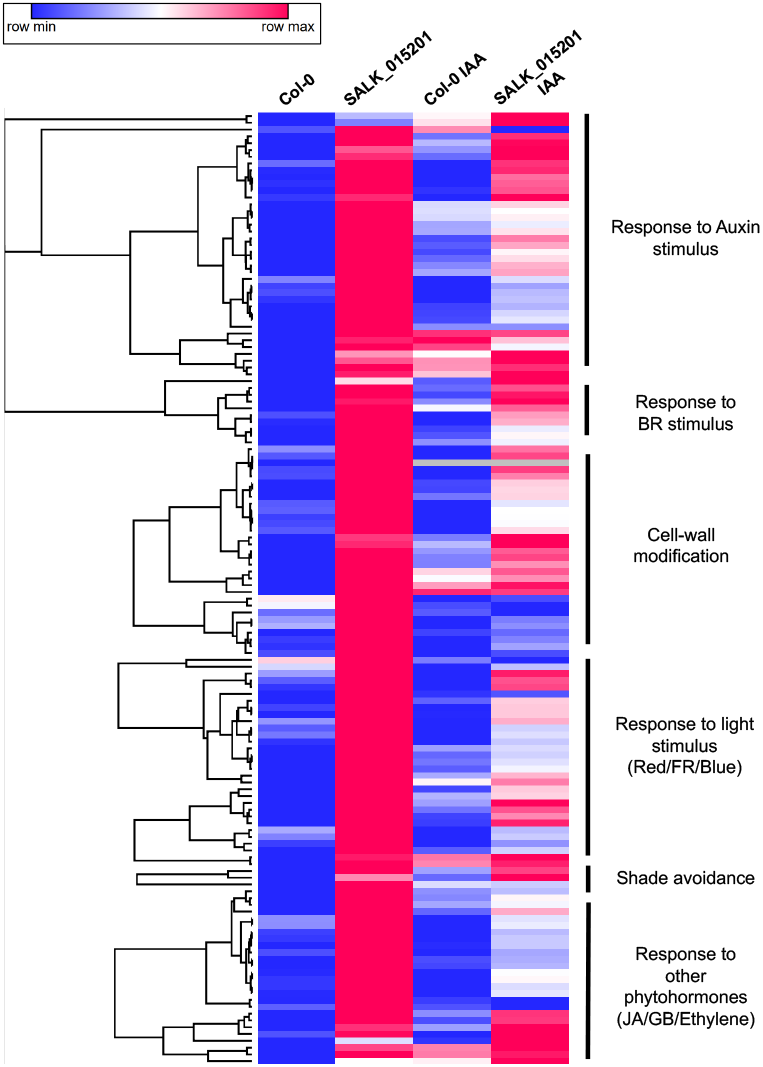
Heatmap of enriched genes in SALK_015201 compared to Col-0 identified by GO analysis. The mean normalized read counts of selected genes with their GO term annotations are shown. The scale bar ranges from lowest (blue) to highest (magenta) relative expression value within a row.

We also observed that multiple transcription factors belonging to the BAP module (i.e. BZR/ARF/PIF families) (Bouré et al., 2019) were upregulated in SALK_015201 as compared to the WT consistent with the elongated hypocotyl phenotype. Classic negative regulators of photomorphogenesis such as BBX28, bHLH transcription factors PIF4, PIF3, PIL1 and PIL6 (Lin et al., 2018; Huq and Quail, 2002; Dong et al., 2017; Luo et al., 2014; Fujimori et al., 2004) showed an increased transcript abundance in SALK_015201 as compared to the WT (Figure 3 and Supplemental Table 1). We also found two of the nine major photoreceptors upregulated in SALK_015201 including the UVA/Blue light sensing PHOT1 (Briggs and Olney, 2001) and far-red light sensing PHYA (Smith, 2000) (Figure 3 and Supplemental Table 1). Our results are congruent with a previous report which discussed the importance of PHYB in photo-regulation of *PHYA* gene expression in plants grown under constant white light or red light (Cantón and Quail, 1999).

Auxin plays a critical role in hypocotyl growth by driving cell expansion (Gray et al., 1998; Zhao et al., 2001). Consistent with this phenomenon we observed numerous auxin inducible genes such as *IAA1, IAA2, IAA7, IAA29, ARF19, ARGOS* to be highly expressed in SALK_015201 compared to WT. Also, genes involved in auxin homeostasis such as *GH3.3, WES1, DFL1* and *NIT2* were upregulated in the mutant. Earlier studies have linked *SMALL AUXIN UP-RNA* (*SAUR*) transcript abundance changes to hypocotyl elongation (McClure and Guilfoyle, 1987; Chae et al., 2012; Spartz et al., 2012; Stamm and Kumar, 2013). Over-expression of *SAUR15* and *SAUR36* promotes hypocotyl elongation because of enhanced cell expansion (Spartz et al., 2014; Stamm and Kumar, 2013). Additionally, *SAUR19, SAUR21*, and *SAUR24* are rapidly induced in the elongating hypocotyls of Arabidopsis seedlings grown in dark (Spartz et al., 2012). We found 18 *SAUR* genes to be elevated in SALK_015201 compared to WT which is consistent with the observed phenotype (Supplemental Table 1). ABCB19, a protein that modulates hypocotyl growth and is required for polar auxin transport (Wu et al., 2016) was also increased in SALK_015201 (Supplemental Table 1). Collectively a large suite of auxin pathway genes were upregulated in SALK_015201 suggesting that altered auxin response and/or metabolism may contribute to the elongated hypocotyl phenotype.

Cell expansion plays a key role in plant growth and development (Braidwood et al., 2014). Cell expansion serves as the fundamental process driving hypocotyl elongation in etiolated Arabidopsis seedlings (Gendreau et al., 1997). EXTENSIN (EXT) and EXPANSIN (EXPs) are two major protein families with known roles in plant cell wall extension and modification (Lamport, 1966; Cosgrove, 2000; McQueen-Mason et al., 1992). We found five *EXP*, four *EXT* and five *XTH* mRNAs to be elevated SALK_015201 which is consistent with the elongated phenotype. The importance of gibberellic acid (GA) as a key phytohormone in regulating the shoot development was determined through genetic and biochemical studies (Koorneef et al., 1985; Jacobsen and Olszewski, 1993; Jacobsen et al., 1996; Peng et al., 1997). Additionally, exogenous application of bioactive GA is known to promote hypocotyl growth in light grown seedlings (Cowling and Harberd, 1999). Congruently, we observed an elevated gibberellin dependent gene expression response in SALK_015201 as compared to the WT (Supplemental Table 1). Genes such as *GA-STIMULATED ARABIDOPSIS 6 (GASA6*) and *GASA14* were found to be elevated in SALK_015201.

### Identification of a background *phyB* mutation, *slim shady*, in SALK_015201

Expression of a *35S:GFP-IRR* transgene in the SALK_015201 background failed to complement the long hypocotyl phenotype (Figure 1C, D), leading us to suspect that SALK_015201 may harbor an unlinked second-site mutation in addition to the annotated T-DNA insertion in *IRR*. Such mutations have been observed in numerous other T-DNA lines (Jiang et al., 2020; Enders et al., 2015; Gao et al., 2015; Yoshida et al., 2018; David et al., 2019) and are thought to be an unintended consequence of the T-DNA integration process (Tax and Vernon, 2001; Castle et al., 1993; Negruk et al., 1996; Nacry et al., 1998; Laufs et al., 1999).

Previously a whole genome sequence was performed for SALK_015201 (Hu et al., 2019). Remapping and variant call analysis of these genomic sequences identified a single base pair deletion of a T nucleotide in the PHYTOCHROME B gene (AT2G18790) at position Chr2:8144002, which corresponds to position 3,370 in the PHYB cDNA (Figure 4A; Supplemental Figure 7). We confirmed the single nucleotide deletion through a derived cleaved amplified polymorphism (dCAPS) assay (Neff et al., 1998) where we introduced an ApoI restriction enzyme site by using primers that contained a single nucleotide mismatch to the template DNA. In this assay, the wild-type *PHYB* allele will digest into 26 and 252 bp while the mutant slim shady allele cannot be cleaved. The SALK_015201 genomic DNA produced a single 277 bp band in this assay while Col-0 and *irr-2* produced the expected wild-type *PHYB* allele PCR products (Figure 4C).

**Fig. 4.**
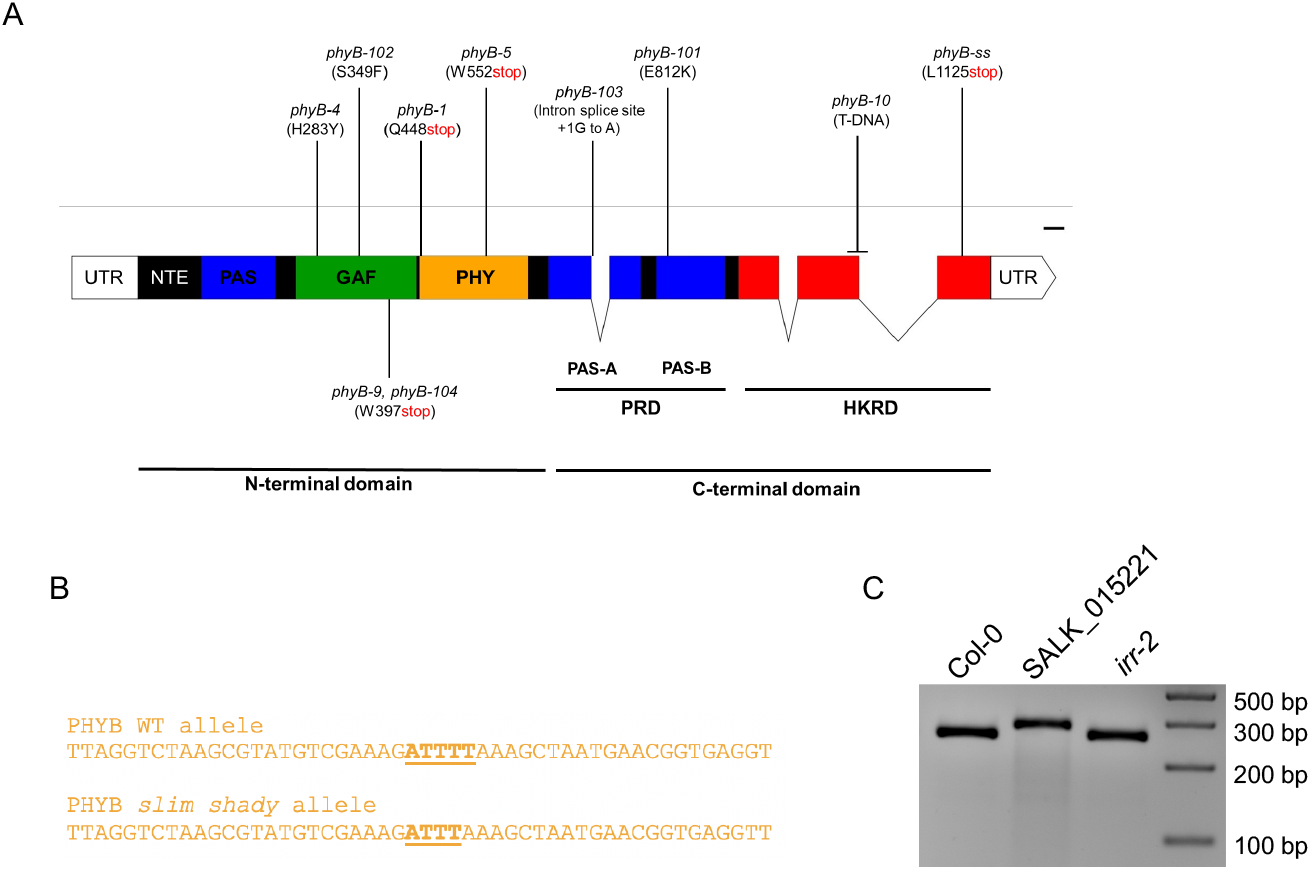
(A) Whole genome sequencing of SALK_015201 indicates a single base pair deletion in *PHYTOCHROME B* at position 3,370 which would lead to a premature stop codon after amino acid L1125 within the Histidine Kinase-Related (HKR) domain. (B) A derived cleaved amplified polymorphic sequence (dCAPS) marker was designed to differentiate between the WT and *phyb-ss* alleles of *PHYB*. (C) Genotyping Col-0, SALK_015201 and *irr-2* lines for *PHYB* and *phyb-ss* alleles using the dCAPS marker. Both Col-0 and *irr-2* are homozygous for the WT *PHYB* allele while SALK_015201 is homozygous for the *phyb-ss* allele.

We also performed PCR and Sanger sequencing reactions on genomic DNA samples from Col-0, SALK_015201 and *irr-2* to confirm this single T deletion within the *PHYB* locus. This analysis confirmed the presence of a single base pair deletion in *PHYB* in the SALK_015201 background that was not observed in Col-0 or *irr-2* (Supplemental Figure 4). This single nucleotide deletion in the *PHYB* coding sequence is predicted to result in a conversion of L1125 to a stop codon in the c-terminal region of the PHYB protein and lead to a truncated protein lacking the last 48 amino acids after I1124 (Figure 4A). The annotated histidine kinase domain of PHYB occurs at amino acid positions 934-1153; thus, the *slim shady* allele of the PHYB would lack a functional kinase domain. The predicted truncated protein would be similar to previously reported C-terminal domain mutations of *phyB* which reduce its biological activity (Qiu et al., 2019; Wagner and Quail, 1995; Wagner et al., 1996; Matsushita et al., 2003).

### Genetic complementation analyses confirm that slim shady is an allele of phyB

The long hypocotyl phenotype we observed in SALK_015201 is due to a loss-of-function allele in *PHYB*. We predicted that *phyb-9* would fail to complement the long hypocotyl phenotype in SALK_015201 (Figure 5A). Conversely, *irr-2* should complement *phyb-9*. We crossed *phyB-9* with both SALK_015201 and *irr-2* and collected F1 seed. F1 seeds and homozygous parental lines were plated on 0.5X LS media supplemented with 1% sucrose to examine hypocotyl phenotypes. The *phyB-9* x *irr-2* F1 plants displayed wild-type hypocotyl phenotypes but all F1 offspring from the *phyB-9* x SALK_015201 crosses exhibited long hypocotyl phenotype (Figure 5B). Thus, *slim shady* represents a novel loss of function allele in *PHYB*.

**Fig. 5.**
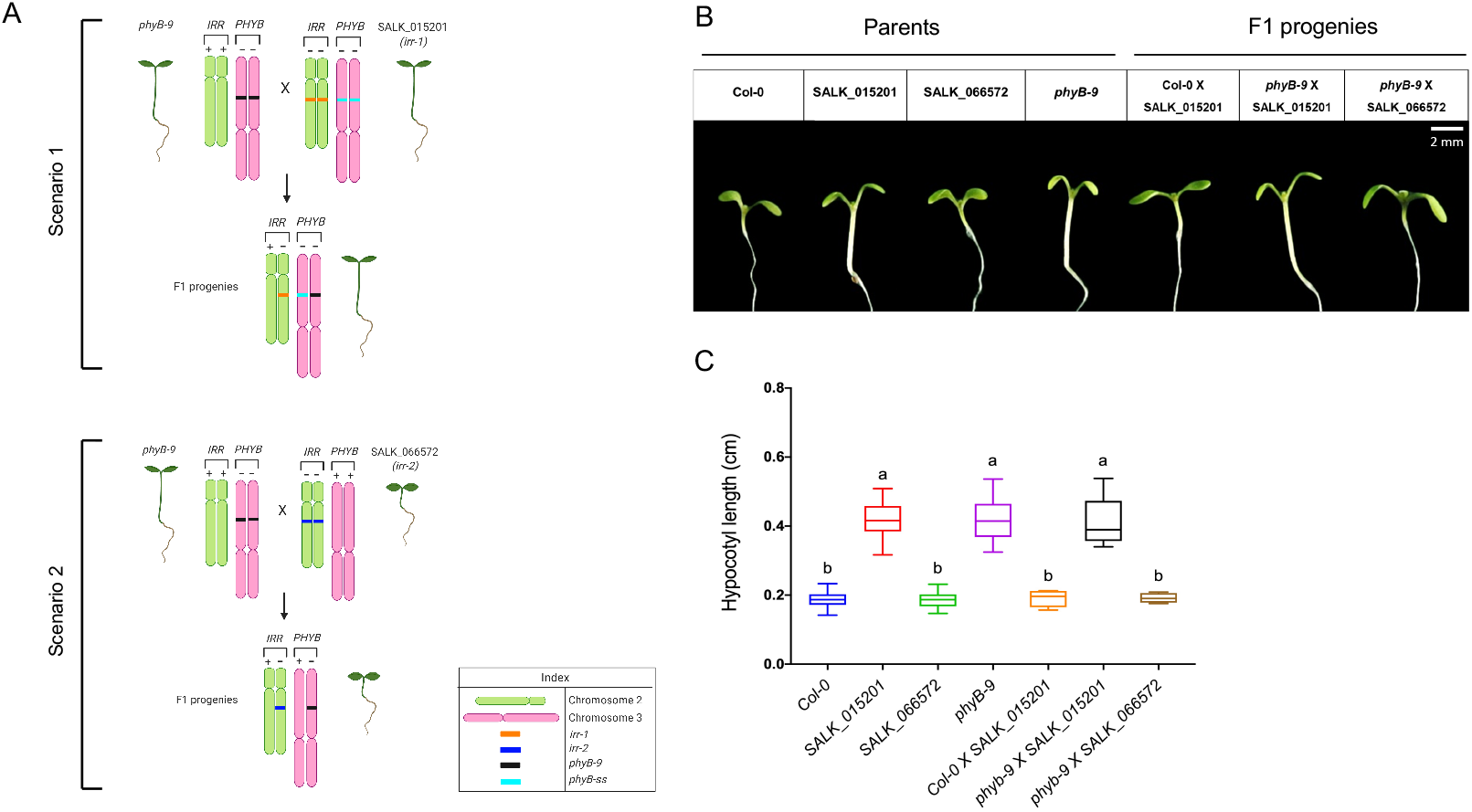
(A) Cartoon schematic of genetic complementation analyses. In scenario 1, the genetic complementation fails because of the *phyB-ss* mutation in SALK_015201 and the resulting F1 offspring have long hypocotyls. In scenario 2, the genetic complementation occurs because the SALK_066572 (*irr-2*) background contains a wildtype *PHYB* allele and the resulting F1 offspring have a wild-type hypocotyl length. (B) Light micrographs of parental and F1 offspring seedlings showing representative hypocotyl lengths. Scale bar = 2 mm. (C) Quantification of hypocotyl lengths following genetic complementation analyses. Letters “a” and “b” indicate the significant statistical differences between the hypocotyl lengths (cm) of different genotypes, determined by a one-way ANOVA followed by Tukey’s HSD post hoc test with p ≤ 0.05. *Eachcolorrepresentsauniquegenotype*.

## Discussion

We recently characterized auxin responsive proteomes (Kelley et al., 2017) and used these data to perform a reverse genetic screen (Page and Grossniklaus, 2002) for seedlings with altered auxin-dependent phenotypes. One of the candidate genes included in this screen was *AT3G23900* which has been recently characterized as *IRR*, a novel RNA-binding protein that is involved in alternatively splicing of its target genes that serve as key defense regulators against pathogen in Arabidopsis (Dressano et al., 2020). Our initial screen was performed with hundreds of SALK T-DNA insertion lines obtained as homozygous so-called SALK “C” lines from the ABRC (O’Malley and Ecker, 2010) including SALK_015201C which contains a knock-out out allele of *IRR*, designated as *irr-1* in (Dressano et al., 2020). This allele shows an elongated hypocotyl phenotype (Figure 1) which has been reported earlier (Petrov et al., 2013). Expressing a *35S:GFP-IRR* in the SALK_015201C background failed to rescue the wild type hypocotyl phenotype (Figure 4), that lead us to suspect the presence of a second-site mutation in this SALK line.

A single base pair deletion in the *PHYB* locus within the last exon of the coding sequence was identified through remapping and variant call analyses of publicly available whole genome sequences from SALK_015201. We further reconfirmed the deletion through a dCAPS assay as well as through Sanger sequencing of the PHYB genomic locus (Figure 4 and Supplemental Figure 4). The missing ‘T’ nucleotide at position 3,370 in the PHYB cDNA, converts a leucine codon to a premature stop codon that likely results in a truncation of 48 amino acids from its C-terminal domain (Figure 4A). Other C-terminal module disruptions to the PHYB protein have been reported to disrupt PHYB activity through various mechanisms (Rockwell et al., 2006; Qiu et al., 2019; Wagner and Quail, 1995).

Previous studies have shown that the dimerization of the PHYB C-terminal domain is essential for its accumulation in subnuclear photobodies as well as its interaction with PIFs driving their photoactivated degradation (Chen et al., 2003; Qiu et al., 2019). The catalytic ATP-binding domain of PHYB contains four conserved subdomains named N, G1, F, and G2 that are essential for its Histidine Kinase Related Domain mediated dimerization (Yeh and Lagarias, 1998; Qiu et al., 2019; Schneider-Poetsch et al., 1991). The slim shady allele of PHYB we have discovered contains an indel that would result in a truncation of the entire G2 subdomain, which would likely generate a loss of function allele. The loss of phyB function is further evident from the long hypocotyl phenotype of *slim shady* that is indistinguishable from *phyB-9* (Figure 1), another well validated knock-out allele of *PHYB* (Reed et al., 1994).

Light activated phytochromes phosphorylate the PIF proteins and target them for 26S proteasomal mediated degradation. PIFs transcriptionally regulate expression of down-stream growth factors and hence pave the mechanism by which phytochromes post-translationally regulate photomorphogenesis (Al-Sady et al., 2006; Park et al., 2012; Hornitschek et al., 2012). Our transcriptomic data show that a subset of ‘photomorphogenic suppressors such as *BBX28, PIF4*, *PIF3*, *PIL1* and *PIL6* are upregulated in *slim shady* as compared to Col-0, that are otherwise tightly regulated. Hence, in addition to degradation of PIFs, photo activated PHYB might be promoting photomorphogenesis through another mechanism in which they suppress the expression of these photomorphogenic suppressors. In future, this multilevel (transcriptional and post-translational) regulation of PIF transcription factors by PHYB and possibly other phytochromes, can be genetically and biochemically explored with the help of non-phosphorylatable PIF transgenic lines.

Transcription factors belonging to BZR, ARF and PIF families, otherwise known as the BAP module, work synergistically to promote the transcription of genes such as *IAA19, SAUR15* and *PRE1*, that are involved in positively regulating hypocotyl growth (Bouré et al., 2019). Our transcriptomic data shows that these downstream growth regulators are collectively elevated in the *slim shady* allele, which can explain the elongated hypocotyl phenotype. In addition, our observation fits with the available knowledge that PHYB suppresses the expression *SAUR15* and *PRE1* by directly interacting with their respective transcriptional activators BES1 and BZR1 and inhibits their promoter binding (Wu et al., 2019). PIF4 competitively inhibits PHYB-BZR1 interaction, therefore possibly attenuating the inhibitory effect of PHYB on the transcriptional activity of BZR1 (Dong et al., 2020). Because these targets are also elevated in *slim shady*, we predict that altered PHYB activity is responsible for the observed altered gene expression patterns.

The uninhibited transcriptional activity of PIFs results in an overaccumulation of auxin followed by a strong auxin response (Ren and Gray, 2015). Auxin induces apoplast acidification through activation of H+-ATPase proton pumps, leading to a cascade of multiple events that in concert contribute towards cell wall expansion, including activation of potassium ion channels and stimulating carbohydrate remodeling proteins such as EXPs and XTHs (Majda and Robert, 2018). Influx of potassium into the cytosol triggers water uptake that builds tensile stress on the cell wall. Simultaneously, proteins like EXPs and XTHs induce cell-wall loosening by disintegrating polysaccharide and glycosidic linkages that support the foundation of plant cell wall. In the case of slim shady, the overexpression of numerous *EXPs (EXPA3, EXPA8, EXPA11, EXLA1* and *EXLA2*) and *XTHs (XTH4, XTH15, XTH17, XTH19, XTH24, XTH30*) are consistent with the observed elongated hypocotyl phenotype.

*SAURs* are a class of early auxin responsive genes that facilitate hypocotyl growth by inducing cell expansion. They are triggered by upstream signals from either of the auxin, BR or light signaling pathways and activate the plasma membrane H+-ATPases by directly binding with and suppressing their inhibitors PP2C-D subfamily of type 2C protein phosphatases (Takahashi et al., 2012; Spartz et al., 2014; Ren et al., 2018). Specifically, over expression of *SAUR15* manifests elongated hypocotyl by facilitating auxin biosynthesis and cell wall expansion via. activation of H+-ATPases in Arabidopsis (Spartz et al., 2014). Transcriptomic data from the novel *PHYB* loss of function allele *slim shady* hints towards a molecular framework for light activated auxin-mediated control of plant cell wall expansion where the loss of PHYB mediated transcriptional inhibition leads to overexpression of genes like *SAUR15, SAUR19, SAUR21* and *SAUR36* that have been implicated for promoting hypocotyl growth (Spartz et al., 2012, 2014; Stamm and Kumar, 2013).

Previous studies have implicated negative photoregulation of *PHYA* expression mediated by PHYB, where *PHYA* gets highly expressed in a *phyB* loss of function background (Cantón and Quail, 1999). As anticipated, the transcript levels of *PHYA* were found to be elevated in *slim shady* as compared to WT. Additionally, PHYA is known to promote hypocotyl growth in light grown Arabidopsis seedlings, caused by the reduction/absence of biologically active PHYB-Pfr (Casal, 1996). Overexpression of PHYA alone has been attributed with the inhibition of hypocotyl growth (Boylan and Quail, 1991). Interestingly, another major UVA/Blue light sensing photoreceptor *PHOT1* showed increased expression in *slim shady*, indicating that PHYB may play a role in *PHOT1* expression in a similar fashion to *PHYA* (Sullivan et al., 2016).

Collectively our initial phenotypic and molecular charac-terization of SALK_015201 suggests that the new *phyB* allele, *slim shady*, found within this T-DNA background may be useful for the photoreceptor and/or auxin community. This study also has implications for functional genomic studies involving T-DNA lines and serves as a reminder that careful genetic analyses require characterization more than one allele, and/or complementation via transgenic approaches to provide conclusive evidence when examining gene function (Bergelson et al., 2016). We encourage all Arabidopsis researchers working with T-DNA lines to confirm target gene expression, obtain more than one T-DNA allele, generate additional alleles via targeted gene editing, or complement with a transgene or BAC when possible in order to confidently ascribe gene function. Original T-DNA collections, many of which were phenotypically screened by numerous laboratories, may contain a wealth of untagged background mutations which could now be identified using next generation sequencing.

## ACKNOWLEDGEMENTS

This work was supported by start-up funds to D.R.K from Iowa State University and NSF grant 1444503 to B.P.D. All seed stocks were obtained from the Arabidopsis Biological Resource Center (ABRC) at Ohio State University.

## Author Contributions

Designed the research: L.D., D.R.K., and B.P. D. Performed research: L.D., L.M., R.E.M., C.M., D.R.K. and J.W.W. Analyzed data: L.D., R.E.M, C.M., B.P.D., D.R.K. Wrote the paper: L.D. and D.R.K with input from all authors.

