## Supplemental Figures for "*slim shady* is a novel allele of PHYTOCHROME B present in the T-DNA line SALK_015201"

**Supplemental Figure 1: qPCR of key marker genes.**

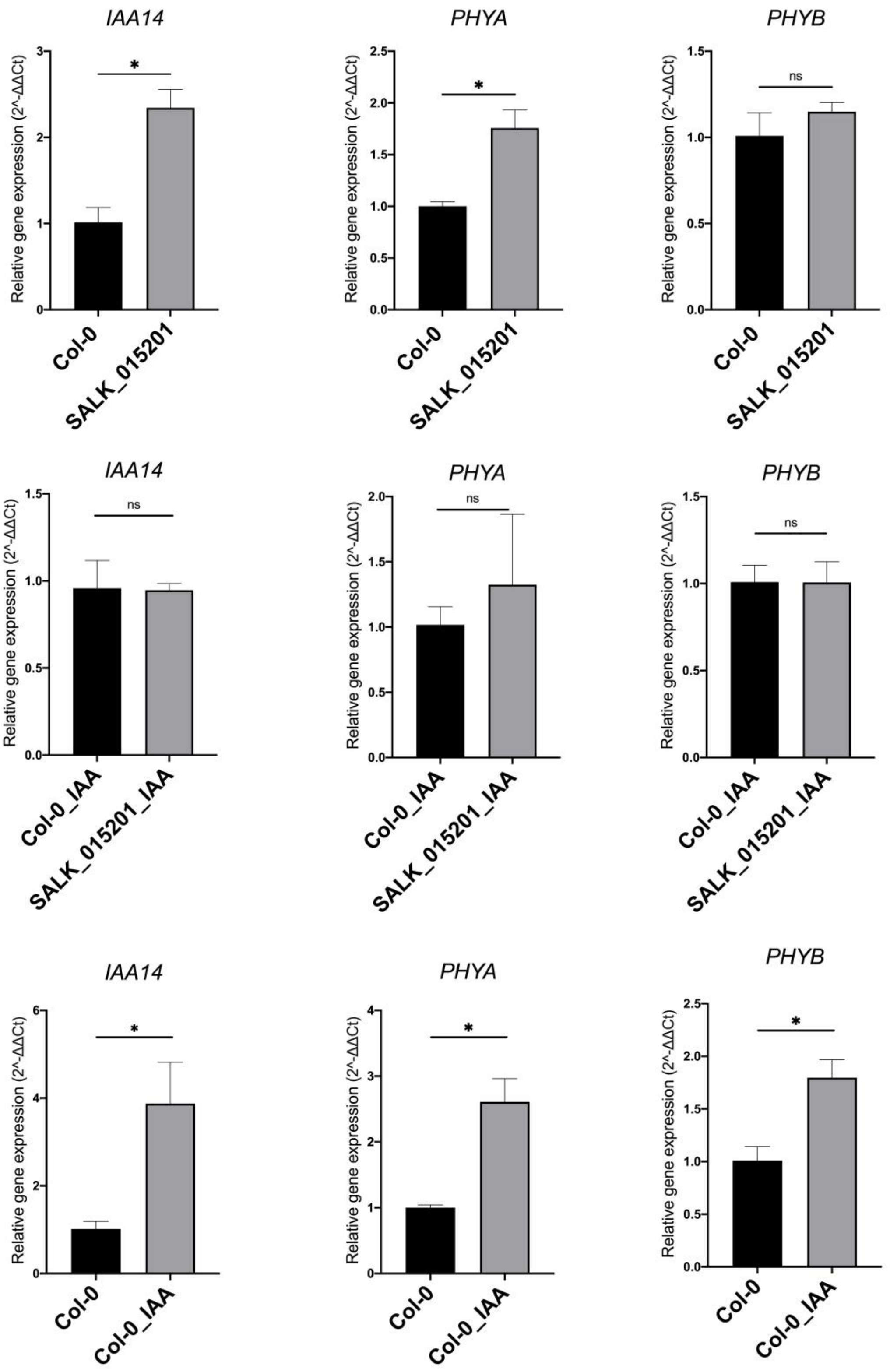

**Supplemental Figure 2: GO network for up-regulated genes in SALK_015201 as compared to Col-0.**

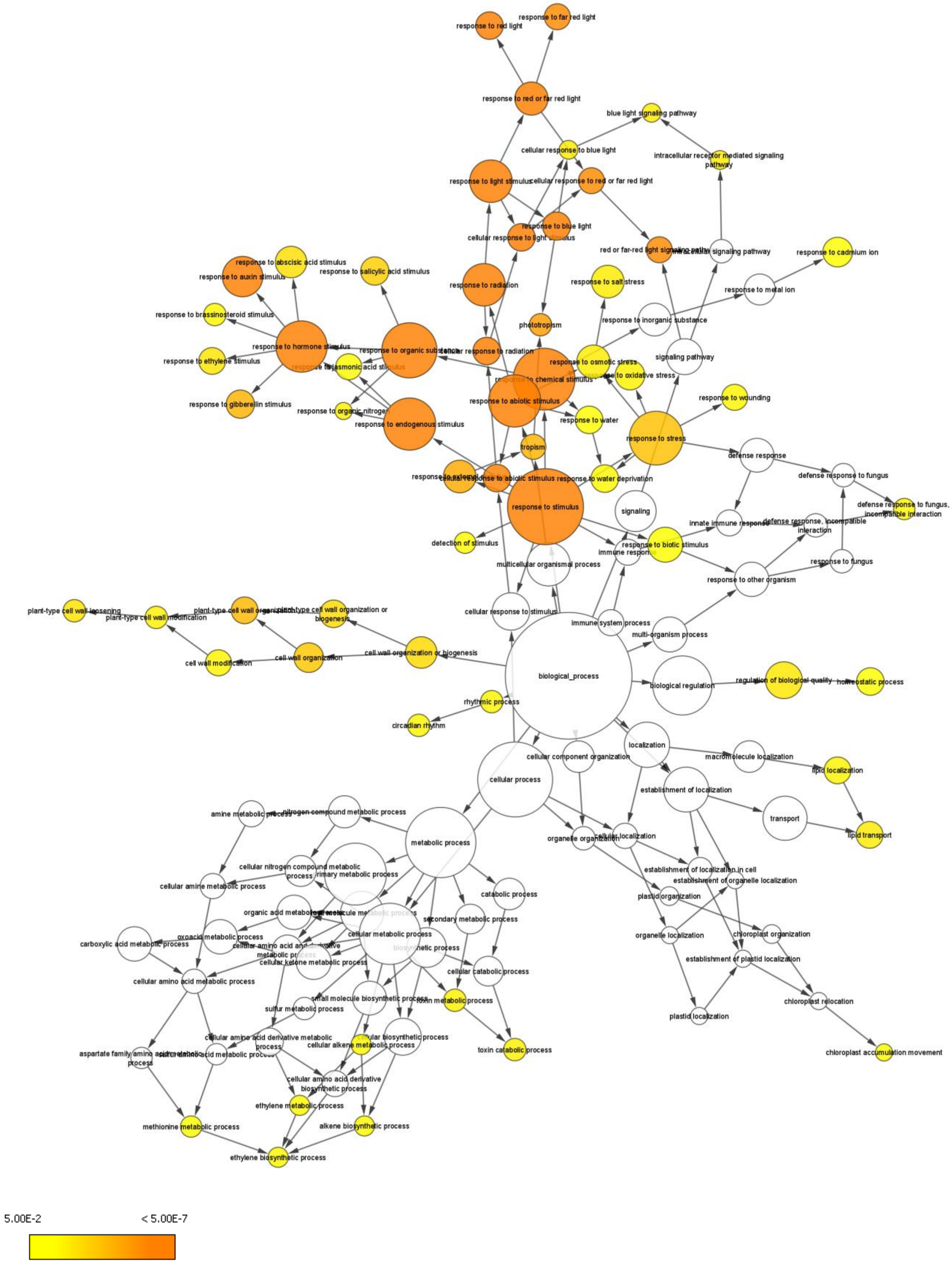

**Supplemental Figure 3: GO network for down-regulated genes in SALK_015201 as compared to Col-0.**

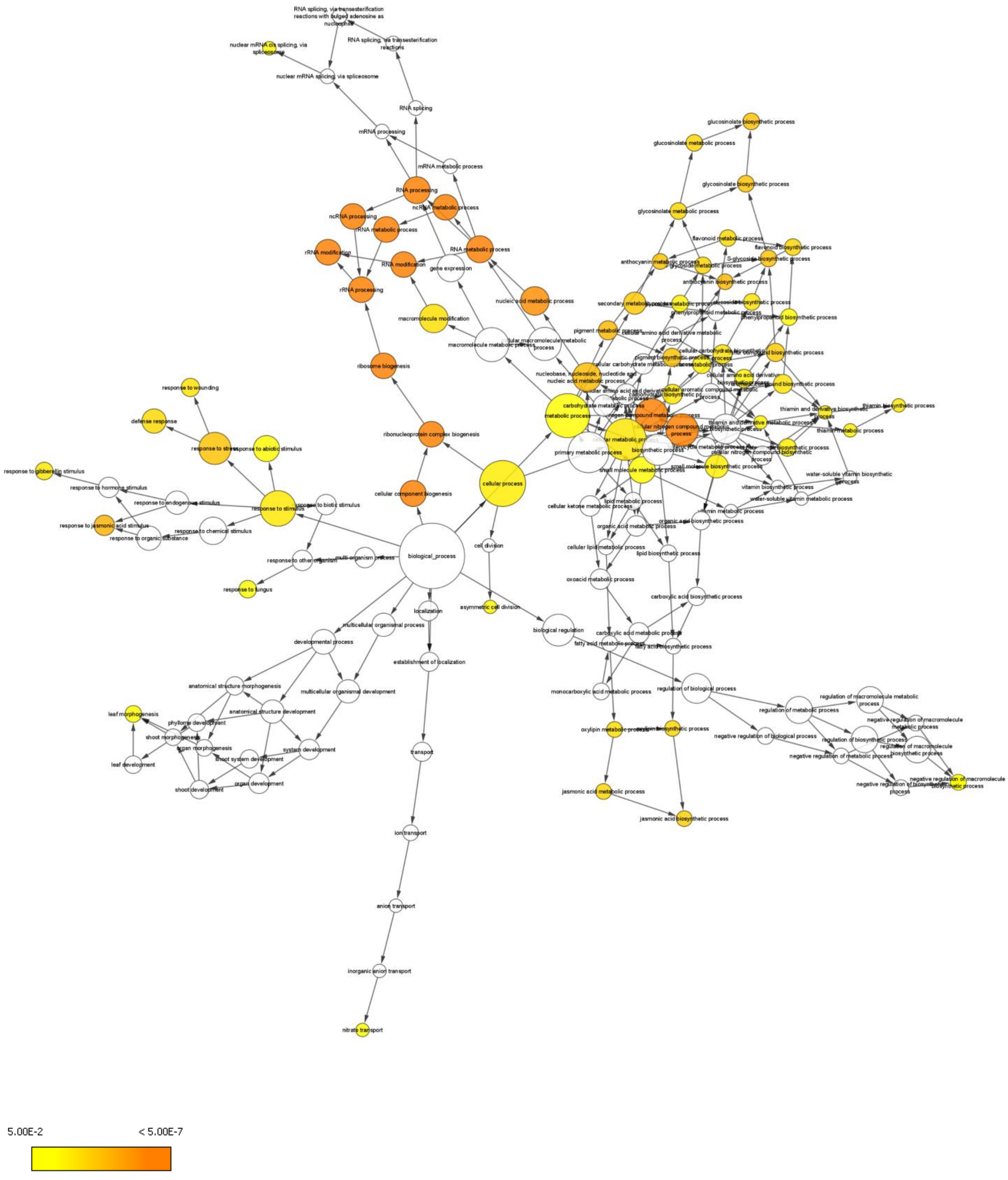

**Supplemental Figure 4: GO network for up-regulated genes in SALK_015201 treated with 1 µM IAA for 24 hours as compared to Col-0 treated with 1 µM IAA for 24 hours.**

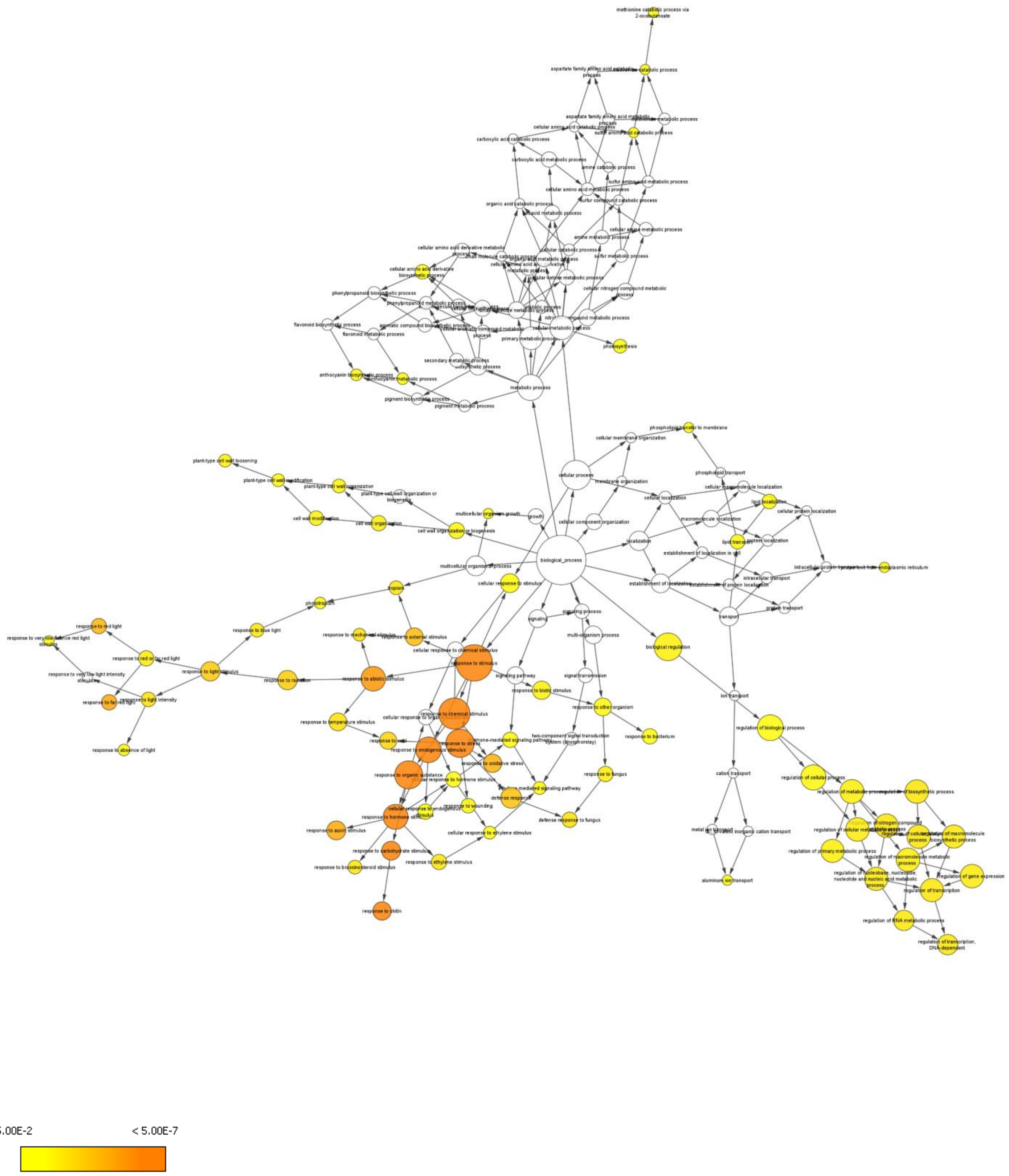

**Supplemental Figure 5: GO network for down-regulated genes in SALK_015201 treated with 1 μM IAA for 24 hours as compared to Col-0 treated with 1 μM IAA for 24 hours.**

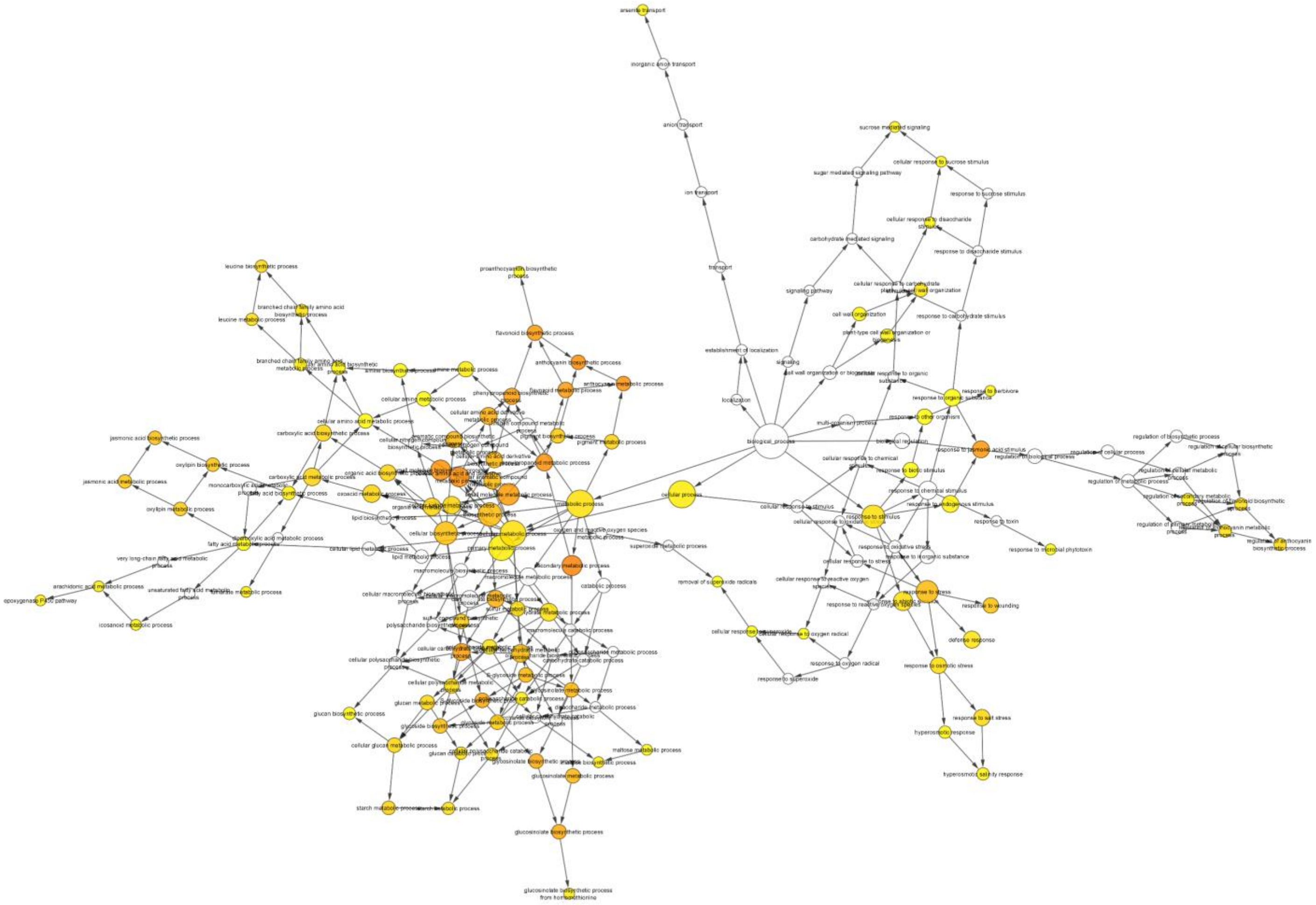

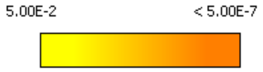

**Supplemental Figure 6: P-value distribution of differentially expressed genes from our 3' RNA seq data set.**

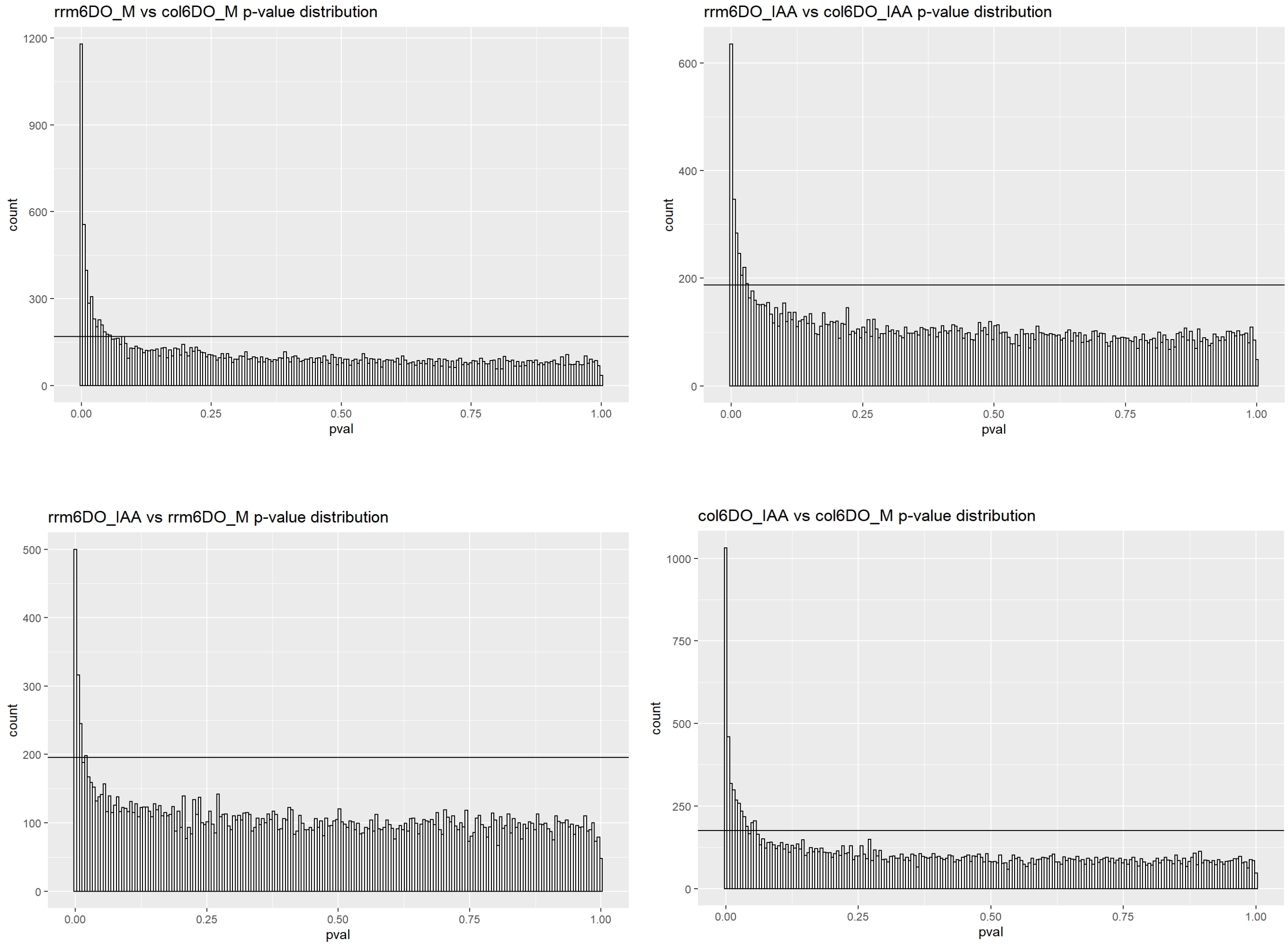

**Supplemental Figure 7: Multiple sequence alignment showing the site of single nucleotide deletion in the *PHYB* locus of SALK_015201 with Col-0 and SALK_066572 as controls and PHYB genomic DNA as reference.**

```
SALK_015201_phyB_F        ------------------------------------------------------------       0
Col-0_phyB_F              ------------------------------------------------------------       0
SALK_066572_phyB_F        ------------------------------------------------------------       0
phyB_genomic_DNA          TTATTTTGTACATAAAGAAAATGAATTTGGTTTGGTTAATTACGAATTTGATTTAGGCGT       3720
SALK_066572_phyB_R        ----------------------------------------------------TTGGGGTT       8
SALK_015201_phyB_R        ---------------------------------------------------TTTGGGGTT       9
Col-0_phyB_R              --------------------------------------------------ATTTTGGGGT       10


SALK_015201_phyB_F        -----------------------ACCATTCCATTTGTTTTGGTTATTGTTTAGTTGGAAC       37
Col-0_phyB_F              -----------------------CCCATTCTATTTGTTTTGGTTATTGTTTAGTTGGAAC       37
SALK_066572_phyB_F        -----------------------CCCATTCTATTTGTTTTGGTTATTGTTTAGTTGGAAC       37
phyB_genomic_DNA          TTAAAGAATTTGAGGTTTTAACCAATTCACTATTTGTTTTGGTTATTGTTTAGTTGGAAC       3780
SALK_066572_phyB_R        TAAAAAGATTTGAGGTTTTAACCAATTCACTATTTGTTTTGGTTATTGTTTAGTTGGAAC       68
SALK_015201_phyB_R        TTAAAAAATTTGAGGTTTTAACCAATTCACTATTTGTTTTGGTTATTGTTTAGTTGGAAC       69
Col-0_phyB_R              TTAAAAGATTTGAGGTTTTAACCAATTCACTATTTGTTTTGGTTATTGTTTAGTTGGAAC       70
                                                       * *****************************

SALK_015201_phyB_F        CTAGATTAGTTTGATTTTTGTATTCGGTTTAGTCGACTTGGGAACTTTTAGACACATCCA       97
Col-0_phyB_F              CTAGATTAGTTTGATTTTTGTATTCGGTTTAGTCGACTTGGGAACTTTTAGACACATCCA       97
SALK_066572_phyB_F        CTAGATTAGTTTGATTTTTGTATTCGGTTTAGTCGACTTGGGAACTTTTAGACACATCCA       97
phyB_genomic_DNA          CTAGATTAGTTTGATTTTTGTATTCGGTTTAGTCGACTTGGGAACTTTTAGACACATCCA       3840
SALK_066572_phyB_R        CTAGATTAGTTTGATTTTTGTATTCGGTTTAGTCGACTTGGGAACTTTTAGACACATCCA       128
SALK_015201_phyB_R        CTAGATTAGTTTGATTTTTGTATTCGGTTTAGTCGACTTGGGAACTTTTAGACACATCCA       129
Col-0_phyB_R              CTAGATTAGTTTGATTTTTGTATTCGGTTTAGTCGACTTGGGAACTTTTAGACACATCCA       130
                          ************************************************************

SALK_015201_phyB_F        TAGGCCTAGAATTAGCAGTCAAGGAATGTAATGTTTTCAAATTGATGAAAACCAGCTCAA       157
Col-0_phyB_F              TAGGCCTAGAATTAGCAGTCAAGGAATGTAATGTTTTCAAATTGATGAAAACCAGCTCAA       157
SALK_066572_phyB_F        TAGGCCTAGAATTAGCAGTCAAGGAATGTAATGTTTTCAAATTGATGAAAACCAGCTCAA       157
phyB_genomic_DNA          TAGGCCTAGAATTAGCAGTCAAGGAATGTAATGTTTTCAAATTGATGAAAACCAGCTCAA       3900
SALK_066572_phyB_R        TAGGCCTAGAATTAGCAGTCAAGGAATGTAATGTTTTCAAATTGATGAAAACCAGCTCAA       188
SALK_015201_phyB_R        TAGGCCTAGAATTAGCAGTCAAGGAATGTAATGTTTTCAAATTGATGAAAACCAGCTCAA       189
Col-0_phyB_R              TAGGCCTAGAATTAGCAGTCAAGGAATGTAATGTTTTCAAATTGATGAAAACCAGCTCAA       190
                          ************************************************************

SALK_015201_phyB_F        AAGTGTAAAACTTGGGTTTCATGTGTTGGTGTCTTTGTTATGTCTTTATTCGTTGTTTGC       217
Col-0_phyB_F              AAGTGTAAAACTTGGGTTTCATGTGTTGGTGTCTTTGTTATGTCTTTATTCGTTGTTTGC       217
SALK_066572_phyB_F        AAGTGTAAAACTTGGGTTTCATGTGTTGGTGTCTTTGTTATGTCTTTATTCGTTGTTTGC       217
phyB_genomic_DNA          AAGTGTAAAACTTGGGTTTCATGTGTTGGTGTCTTTGTTATGTCTTTATTCGTTGTTTGC       3960
SALK_066572_phyB_R        AAGTGTAAAACTTGGGTTTCATGTGTTGGTGTCTTTGTTATGTCTTTATTCGTTGTTTGC       248
SALK_015201_phyB_R        AAGTGTAAAACTTGGGTTTCATGTGTTGGTGTCTTTGTTATGTCTTTATTCGTTGTTTGC       249
Col-0_phyB_R              AAGTGTAAAACTTGGGTTTCATGTGTTGGTGTCTTTGTTATGTCTTTATTCGTTGTTTGC       250
                          ************************************************************

SALK_015201_phyB_F        AGAATGGCGTGTCCAGGTGAAGGTCTGCCTCCAGAGCTAGTCCGAGACATGTTCCATAGC       277
Col-0_phyB_F              AGAATGGCGTGTCCAGGTGAAGGTCTGCCTCCAGAGCTAGTCCGAGACATGTTCCATAGC       277
SALK_066572_phyB_F        AGAATGGCGTGTCCAGGTGAAGGTCTGCCTCCAGAGCTAGTCCGAGACATGTTCCATAGC       277
phyB_genomic_DNA          AGAATGGCGTGTCCAGGTGAAGGTCTGCCTCCAGAGCTAGTCCGAGACATGTTCCATAGC       4020
SALK_066572_phyB_R        AGAATGGCGTGTCCAGGTGAAGGTCTGCCTCCAGAGCTAGTCCGAGACATGTTCCATAGC       308
SALK_015201_phyB_R        AGAATGGCGTGTCCAGGTGAAGGTCTGCCTCCAGAGCTAGTCCGAGACATGTTCCATAGC       309
Col-0_phyB_R              AGAATGGCGTGTCCAGGTGAAGGTCTGCCTCCAGAGCTAGTCCGAGACATGTTCCATAGC       310
                          ************************************************************

SALK_015201_phyB_F        AGCAGGTGGACAAGCCCTGAAGGTTTAGGTCTAAGCGTATGTCGAAAGATT-TAAAGCTA       336
Col-0_phyB_F              AGCAGGTGGACAAGCCCTGAAGGTTTAGGTCTAAGCGTATGTCGAAAGATTTTAAAGCTA       337
SALK_066572_phyB_F        AGCAGGTGGACAAGCCCTGAAGGTTTAGGTCTAAGCGTATGTCGAAAGATTTTAAAGCTA       337
phyB_genomic_DNA          AGCAGGTGGACAAGCCCTGAAGGTTTAGGTCTAAGCGTATGTCGAAAGATTTTAAAGCTA       4080
SALK_066572_phyB_R        AGCAGGTGGACAAGCCCTGAAGGTTTAGGTCTAAGCGTATGTCGAAAGATTTTAAAGCTA       368
SALK_015201_phyB_R        AGCAGGTGGACAAGCCCTGAAGGTTTAGGTCTAAGCGTATGTCGAAAGATT-TAAAGCTA       368
Col-0_phyB_R              AGCAGGTGGACAAGCCCTGAAGGTTTAGGTCTAAGCGTATGTCGAAAGATTTTAAAGCTA       370
                          *************************************************** ********
```

```
SALK_015201_phyB_F      ATGAACGGTGAGGTTCAATACATCCGAGAATCAGAACGGTCCTATTTCCTCATCATTCTG      396
Col-0_phyB_F            ATGAACGGTGAGGTTCAATACATCCGAGAATCAGAACGGTCCTATTTCCTCATCATTCTG      397
SALK_066572_phyB_F      ATGAACGGTGAGGTTCAATACATCCGAGAATCAGAACGGTCCTATTTCCTCATCATTCTG      397
phyB_genomic_DNA        ATGAACGGTGAGGTTCAATACATCCGAGAATCAGAACGGTCCTATTTCCTCATCATTCTG      4140
SALK_066572_phyB_R      ATGAACGGTGAGGTTCAATACATCCGAGAATCAGAACGGTCCTATTTCCTCATCATTCTG      428
SALK_015201_phyB_R      ATGAACGGTGAGGTTCAATACATCCGAGAATCAGAACGGTCCTATTTCCTCATCATTCTG      428
Col-0_phyB_R            ATGAACGGTGAGGTTCAATACATCCGAGAATCAGAACGGTCCTATTTCCTCATCATTCTG      430
                        ************************************************************

SALK_015201_phyB_F      GAACTCCCTGTACCTCGAAAGCGACCATTGTCAACTGCTAGTGGAAGTGGTGACATGATG      456
Col-0_phyB_F            GAACTCCCTGTACCTCGAAAGCGACCATTGTCAACTGCTAGTGGAAGTGGTGACATGATG      457
SALK_066572_phyB_F      GAACTCCCTGTACCTCGAAAGCGACCATTGTCAACTGCTAGTGGAAGTGGTGACATGATG      457
phyB_genomic_DNA        GAACTCCCTGTACCTCGAAAGCGACCATTGTCAACTGCTAGTGGAAGTGGTGACATGATG      4200
SALK_066572_phyB_R      GAACTCCCTGTACCTCGAAAGCGACCATTGTCAACTGCTAGTGGAAGTGGTGACATGATG      488
SALK_015201_phyB_R      GAACTCCCTGTACCTCGAAAGCGACCATTGTCAACTGCTAGTGGAAGTGGTGACATGATG      488
Col-0_phyB_R            GAACTCCCTGTACCTCGAAAGCGACCATTGTCAACTGCTAGTGGAAGTGGTGACATGATG      490
                        ************************************************************

SALK_015201_phyB_F      CTGATGATGCCATATTAGTCACACTTCAGTTGGTATGAGAGTTTGTATCATTGTATGAGT      516
Col-0_phyB_F            CTGATGATGCCATATTAGTCACACTTCAGTTGGTATGAGAGTTTGTATCATTGTATGAGT      517
SALK_066572_phyB_F      CTGATGATGCCATATTAGTCACACTTCAGTTGGTATGAGAGTTTGTATCATTGTATGAGT      517
phyB_genomic_DNA        CTGATGATGCCATATTAGTCACACTTCAGTTGGTATGAGAGTTTGTATCATTGTATGAGT      4260
SALK_066572_phyB_R      CTGATGATGCCATATTAGTCACACTTCAGTTGGTATGAGAGTTTGTATCATTGTATGAGT      548
SALK_015201_phyB_R      CTGATGATGCCATATTAGTCACACTTCAGTTGGTATGAGAGTTTGTATCATTGTATGAGT      548
Col-0_phyB_R            CTGATGATGCCATATTAGTCACACTTCAGTTGGTATGAGAGTTTGTATCATTGTATGAGT      550
                        ************************************************************

SALK_015201_phyB_F      GTTTGTGTGTCTAACGACGTCGGAGGAGGATAGAAAGTTTTTTTTTTGTTTCCGGTGAGA      576
Col-0_phyB_F            GTTTGTGTGTCTAACGACGTCGGAGGAGGATAGAAAGTTTTTTTTTTGTTTCCGGTGAGA      577
SALK_066572_phyB_F      GTTTGTGTGTCTAACGACGTCGGAGGAGGATAGAAAGTTTTTTTTTTGTTTCCGGTGAGA      577
phyB_genomic_DNA        GTTTGTGTGTCTAACGACGTCGGAGGAGGATAGAAAGTTTTTTTTTTGTTTCCGGTGAGA      4320
SALK_066572_phyB_R      GTTTGTGTGTCTAACGACGTCGGAGGAGGATAGAAAGTTTTTTTTTTGTTTCCCGGGAGA      608
SALK_015201_phyB_R      GTTTGTGTGTCTAACGACGTCGGAGGAGGATAGAAAGTTTTTTTTTTTTTCCCGGGAGAT      608
Col-0_phyB_R            GTTTGTGTGTCTAACGACGTCGGAGGAGGATAGAAAGTTTTTTTTTTTTTCCCGGGAGAT      610
                        *********************************************** ** ** *

SALK_015201_phyB_F      TTAGTAGAGAAGAGGGAGATTATTTGCTTCCGCCNNNNNAA-------------------      617
Col-0_phyB_F            TTAGTAGAGAAGAGGGAGATTATTTGCTTCCGCCTTCAGCAA------------------      619
SALK_066572_phyB_F      TTAGTAGAGAAGAGGGAGATTATTTGCTTCCGGNNNCNNNAAA-----------------      620
phyB_genomic_DNA        TTAGTAGAGAAGAGGGAGATTATTTGCGTTCAGCTCAGCTCGCCGGAAAAAAAACGTAAC      4380
SALK_066572_phyB_R      TAGAGAAGGG--------------------------------------------------      618
SALK_015201_phyB_R      TGAGAAGGG---------------------------------------------------      617
Col-0_phyB_R            TGGGAAGGG---------------------------------------------------      619
                        *    *  *
```
